# Adsorption of semiflexible polymers in crowded environments

**DOI:** 10.1101/2021.04.22.440914

**Authors:** Gaurav Chauhan, Michael L. Simpson, Steven M. Abel

## Abstract

Macromolecular crowding is a feature of cellular and cell-free systems that, through depletion effects, can impact the interactions of semiflexible biopolymers with surfaces. In this work, we use computer simulations to study crowding-induced adsorption of semiflexible polymers on otherwise repulsive surfaces. Crowding particles are modeled explicitly, and we investigate the interplay between the bending stiffness of the polymer and the volume fraction and size of crowding particles. Adsorption is promoted by stiffer polymers, smaller crowding particles, and larger volume fractions of crowders. We characterize transitions from non-adsorbed to partially and strongly adsorbed states as a function of the bending stiffness. The crowding-induced transitions occur at smaller values of the bending stiffness as the volume fraction of crowders increases. Concomitant effects on the size and shape of the polymer are reflected by crowding- and stiffness-dependent changes to the radius of gyration. We also demonstrate that curvature of the confining surface can induce desorption when the bending stiffness is sufficiently large. The results of our simulations shed light on the interplay of crowding and bending stiffness on the spatial organization of biopolymers in encapsulated cellular and cell-free systems.

## I. INTRODUCTION

Asakura and Oosawa’s seminal paper on depletion interactions, published in 1954, describes how attractive depletion forces arise between objects due to the presence of smaller solute particles.^1^ Depletion interactions have been studied widely in the context of colloid-polymer and other soft matter systems, and many outstanding questions remain to be addressed.^2,3^ More recently, there has been growing interest in the roles of depletion interactions in biological systems, where they have been suggested to play a role in cellular organization,^4^ genome organization,^5–8^ gene regulation,^9–11^ and controlling intracellular phase separation.^12,13^

The interiors of cells contain large concentrations of macromolecules that can occupy up to 40% of the total cellular volume.^14^ The cellular environment is replete with semiflexible polymers such as DNA, actin, microtubules, etc., which have a wide range of persistence lengths. These polymers are often in the presence of surfaces such as the plasma or nuclear membrane. The abundance of macromolecules can result in attractive depletion interactions between biopolymers and surfaces,^4^ which can lead to adsorption of the polymers. Understanding the effects of crowding on properties of semi-flexible polymers and their interactions with surfaces will help shed light on the role of depletion interactions in organizing cellular systems.

Crowding-induced depletion interactions have been shown to impact the conformations of both flexible and semiflexible biopolymers. For flexible polymers, crowding can cause a coil-to-globule transition,^15^ induce collapse of model chromosomes,^16^ and lead to attraction between ring polymers.^17^ For semiflexible polymers, which have an inherent bending stiffness, experiments have shown that actin filaments undergo condensation upon addition of non-adsorbing polymeric crowders.^18^ Simulations have also shown that polydispersity of crowder sizes and shapes can unexpectedly swell polymers of intermediate bending stiffness.^19^

Crowding can also lead to the adsorption of biopolymers onto surfaces. DNA plasmids have been shown to preferentially localize near the walls of crowded cell-sized vesicles,^20,21^ and by tuning the concentration of depletants, Welch et al. observed actin filaments in both partially and fully adsorbed states.^22^ Interactions of proteins with surfaces in crowded environments have also been shown to enhance the formation of protein fibrils.^23,24^ For flexible ring polymers, the magnitude of crowding-induced attraction to a wall was shown to be notably stronger than the magnitude of polymer-polymer attraction, and an effective polymer-wall attraction emerged at a lower volume fraction of crowding particles.^17^

Understanding interactions of polymers with surfaces has been a long-standing problem of interest in polymer physics.^25–27^ While polymer adsorption has been studied extensively for flexible polymers,^28–20^ much less is known about the adsorption of semiflexible polymers.^27,31^ To characterize adsorption, theories and simulations must account for internal degrees of freedom of the polymer, and stiffer polymers lose less conformational entropy upon adsorption. As a consequence, when there is explicit attraction between a semiflexible polymer and a wall, adsorption is promoted by a larger bending stiffness and the critical strength of attraction required for adsorption decreases with increasing stiffness.^22,31–33^ Milchev and Binder^31^ recently showed that the conformations of partially adsorbed chains are not well-described by the wormlike chain model and that adsorption does not lead to the expected 2-dimensional decay of the orientational correlation function over an intermediate but broad range of the bending stiffness. Additionally, for semiflexible polymers, the curvature of the surface can impact the adsorption behavior due to an energetic bending penalty experienced by polymers when adsorbed to a curved surface.^34–37^

In this paper, we use computer simulations to study crowding-induced interactions of linear semiflexible polymers with surfaces. Previous works studying the adsorption of semiflexible polymers have typically used generic, short-ranged attractive potentials between polymer beads and a surface. In contrast with these works, we explicitly simulate crowding particles and their effect on a bead-spring model of a semiflexible polymer. This allows us to capture features missing from simpler approaches, including (i) details of crowding-induced interactions between polymer beads and walls that may not be reflected in simple potentials and (ii) crowding-induced interactions between different segments of the same polymer. Additionally, it is challenging to determine a quantitative relation between the properties of the crowders (size, volume fraction, etc.) and the strength of the depletion interaction, especially at higher concentrations where the depletion interaction may not scale linearly with the concentration of crowders.

In the following, we explore how polymer adsorption is impacted by the bending stiffness of the polymers and the volume fraction and size of the crowders. We characterize properties of the polymers and show that the shape of the system and curvature of the surface can impact the adsorption. Taken together, our work sheds light on the role of crowding in the spatial organization of semiflexible polymers in cellular and cell-free environments.

## II. METHODS

We studied effects of crowding on a semiflexible polymer in the presence of a surface using Langevin dynamics simulations. The polymer was modeled as a linear, semiflexible chain consisting of *N* = 50 beads. Adjacent beads were connected via the finitely extensible nonlinear elastic (FENE) bond potential,

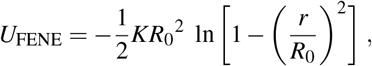

where *r* is the center-to-center distance between two adjacent beads. The diameter of each bead was σ_*m*_ = σ, the maximum distance between two connected beads was *R*_0_ = 1.5σ, and the spring constant was *K* = 15*ε*/σ^2^. σ and *ε* set the length and energy units, and we report lengths and energies in terms of them. The persistence length of the polymer was controlled by changing the bending stiffness, *κ*, of the bending potential,

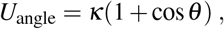

where *θ* is the angle formed by three consecutive beads of the semiflexible polymer. The persistence length (*l_p_*) denotes the length scale over which tangent correlations decay along the polymer chain. The theoretical relationship between the bending stiffness and the persistence length in *d* spatial dimensions is *l_p_* = 2*κ*/(*d* – 1).^34,38,39^

Crowding particles (“crowders”) were modeled as purely repulsive particles of diameter σ_*c*_. We considered two sizes, σ_*c*_ = 0.8σ and σ. The volume fraction of crowders, *ϕ* = *N_c_*(4/3)*π*(σ_*c*_/2)^3^/*V*_box_, was controlled by changing the number of crowder particles in the simulation box of volume *V*_box_. All particles (polymer beads and crowders) interacted via the short-ranged and purely repulsive Weeks-Chandler-Andersen (WCA) potential,^40^

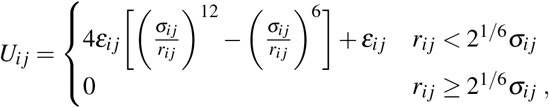

where *r_ij_* is the center-to-center distance between particles *i* and *j*. The strength parameter was the same for all pairs, *ε_ij_* = *ε* = *k*_B_*T*. Further, σ_*ij*_ = (σ_*i*_ + σ_*j*_)/2, where σ_*i*_ and σ_*j*_ denote the diameter of particles *i* and *j*, respectively.

We simulated a single polymer with many crowders in the presence of confining surfaces, which were represented by the purely repulsive 9-3 Lennard-Jones potential,

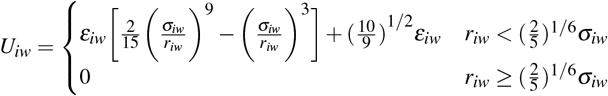

Here, *r_iw_* is the closest distance between the particle and the wall, the strength factor for wall-particle interactions was the same for all particles (*ε_iw_* = *ε* = *k*_B_*T*), and the length parameter σ_*iw*_ was specified by (2/5)^1/6^ σ_*iw*_ = σ_*ii*_/2. To simulate a flat wall, we used a repulsive wall in the *z*-direction with periodic boundaries in the *x*- and *y*-directions. The dimensions of the simulation box were 50σ in the *x*- and *y*-dimensions and 40σ in the *z*-dimension. We also simulated polymers in spherical confinement with a radius of 30σ using the above potential.

The Langevin equation was integrated forward in time using the velocity-Verlet algorithm in the LAMMPS simulation package.^41,42^ The timestep for integration was 0.005*τ*, where *τ* is the natural unit of time. The total number of timesteps was typically 4.95 × 10^7^, with data sampled after an initial equilibration time of 2 × 10^7^ timesteps. For systems at large crowding fractions (*ϕ* = 0.4), we ran simulations for an additional 5 × 10^7^ timesteps, for a total sampling time of 7.95 × 10^7^ timesteps. Averages of dynamical variables were calculated by time averaging along each trajectory. Resulting trajectories were visualized using OVITO.^43^

## III. RESULTS

In cellular and cell-free systems, the linear size of macromolecular crowders is often comparable to the cross-sectional diameter of biopolymers such as DNA or actin. Widely-used synthetic crowders such as dextran 70 and Ficoll 70 have hydrodynamic radii of ≈ 6.8 nm and ≈ 5 nm, respectively.^44^ These sizes, which can be varied with molecular weight, are similar to the effective diameters of actin (≈ 6 nm)^22^ and ds-DNA (≈ 6.6 nm at 60 mM ionic strength).^45^ As such, we simulated crowder sizes that were comparable to the size of a bead of the polymer chain.

### A. Interplay between polymer stiffness, volume fraction of crowders, and crowder size

As a measure of association of the polymer with the wall, we characterized the number of polymer beads within distance σ of a wall (*N*_wall_). Figure 1 shows 〈*N*_wall_/*N*〉, the average fraction of polymer beads in close proximity to a wall, as a function of bending stiffness (*κ*) for various crowding fractions (*ϕ*). When *ϕ* = 0, there are no depletion interactions, and the polymer interacts with the walls only via short-ranged repulsive interactions between its beads and the wall. In this regime, 〈*N*_wall_/*N*〉 remains close to zero for all *κ*.

When crowders were present, we first considered the case in which the crowding particles were the same size as the monomer beads (σ_*c*_ = σ_*m*_, dashed lines in Fig. 1). For a flexible chain (*κ* = 0), the average fraction of monomers near the wall remained close to zero for all crowding fractions up to *ϕ* = 0.4. Thus, the flexible chains did not experience an appreciable crowding-induced attraction to the wall. Physically, there is a reduction in the conformational entropy of the polymer when it is close to the wall. In the absence of crowders, this leads to an effective repulsion between the center of mass of the polymer and the wall as well as a reduction in the density of monomers near the wall.^17^ Our results indicate that when *κ* = 0 and σ_*c*_ = σ_*m*_, depletion effects do not overcome the loss of entropy resulting from the association of the polymer with a wall. That is, when the polymer is in close contact with the wall, the entropic gains by crowding particles due to a larger effective volume do not offset the loss of conformational entropy of the polymer.

Even though there was no crowding-induced adsorption for flexible polymers, we observed that increasing the bending stiffness resulted in association with a wall when *ϕ* ≥ 0.2. For *ϕ* = 0.2, weak adsorption emerged at *κ* = 50 and 100. In this regime, the adsorption is characterized by partial contact with the wall. The contact is transient, being characterized by intermittent excursions away from the wall. For *ϕ* = 0.3 and 0.4, we observed a transition to strong adsorption 〈*N*_wall_/*N*〉 ≈ 1) with increasing *κ*. The transition occurs at smaller values of *κ* for the larger crowding fraction. When the polymer was strongly adsorbed, we did not observe excursions of the center of mass away from the wall during the timescale of the simulations.

**FIG. 1.**
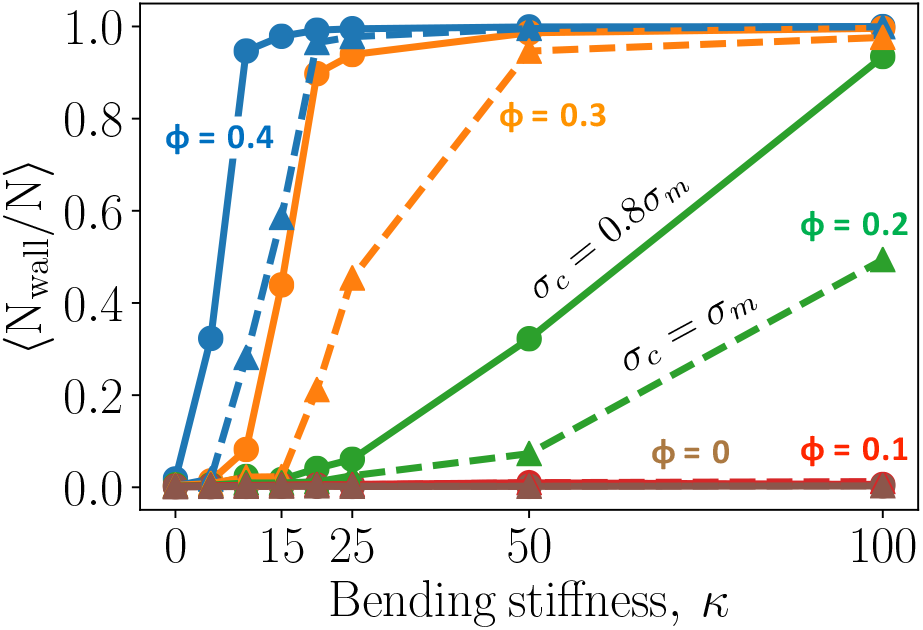
Average fraction of monomers near a wall, 〈*N*_wall_/*N*〉, as a function of the bending stiffness, *κ*. Two sizes of the crowding particles are shown: σ_*c*_ = σ_*m*_ (dashed) and σ_*c*_ = 0.8σ_*m*_ (solid). Different volume fractions of the crowding particles (*ϕ*) are shown.

Previous work has shown that smaller crowding particles can induce considerable compaction of flexible polymers^15^ and strong attraction of a flexible ring polymer to a wall.^17^ We studied the effect of crowder size on polymer adsorption by considering smaller crowding particles with σ_*c*_ = 0.8σ_*m*_ (Fig. 1, solid lines). Note that the volume of each particle is roughly half that of the previous case. At the same values of *ϕ* and *κ*, smaller crowding particles promote stronger adsorption. This is reflected by (i) the transition to strong adsorption (〈*N*_wall_/*N*〉 ≈ 1) at smaller values of *κ* and (ii) the larger fraction of monomers close to a wall in the weak and moderate adsorption regimes. Thus, the strength of the depletion interactions increased with a decrease in crowder size at fixed volume fraction, which is consistent with previous work.^15,17^ In the rest of the paper, we focus on the smaller crowding particle with σ_*c*_ = 0. 8σ_*m*_.

Figure 2 further explores the contact of the polymer with the wall. It shows the full distribution of contact fractions for different *ϕ* and *κ*. There is a prominent peak at *N*_wall_/*N* = 0 for cases without adsorption (e.g., *ϕ* = 0.2, *κ* = 0), indicating that the polymer rarely contacts the wall in these cases. Analysis of trajectories with *ϕ* ≤ 0.1 produced similar results with a prominent peak at 0. For *ϕ* = 0.2, adsorption occurs at sufficiently large values of *κ*. In Fig. 2, the first hint of this transition can be seen at *κ* = 15, where the peak at *N*_wall_/*N* = 0 is slightly lower than for smaller values of *κ*, indicating a slight increase in the weight of configurations in which the polymer is in contact with a wall. For *κ* = 50, this feature is more prominent, and a broad distribution of contact fractions can be observed, including configurations in which the polymer is fully in contact. This is reflective of partial adsorption, in which fluctuations are prominent: The polymer is commonly not associated with the wall, but when it is, it exhibits a broad distribution of the number of monomers in contact. Similar behavior is observed at *ϕ* = 0.3 with *κ* = 15 and at *ϕ* = 0.4 with *κ* = 5, also indicating weak, partial adsorption. For crowding fractions of *ϕ* = 0.3 and 0.4, the probability distribution is peaked at *N*_wall_/*N* = 1 for the stiffest polymers, indicating strong adsorption in which the polymer is typically in complete contact with the wall.

### B. Properties of the polymers

The distributions of weakly adsorbed polymers in Fig. 2 indicate a broad distribution of the fraction of monomers in contact with the wall. Figure 3 shows representative snapshots of the polymer in the partially adsorbed states. Conformations of partially adsorbed polymers are often described using the terminology of trains, tails, and loops.^22^ Trains are continuous adsorbed segments of the polymer, tails are non-adsorbed end segments of the polymer, and loops are non-adsorbed segments between two trains. Examples for *ϕ* = 0.2, 0.3, and 0.4 are shown; as the crowding fraction increases, a smaller bending stiffness is needed for the polymer to adsorb to the wall. Three snapshots are shown for *ϕ* = 0.2 and *κ* = 50. In each, there is a single train along with one or two tails. Because the polymer is relatively stiff, it would be energetically costly for a loop to form (due to high local curvature). Hence, monomers not in contact with the wall tend to be located at the tails of the polymer. As the crowding fraction increases, the presence of tails decreases and eventually, in the limit of strong adsorption, the polymer is completely in contact with the wall as a single train.

**FIG. 2.**
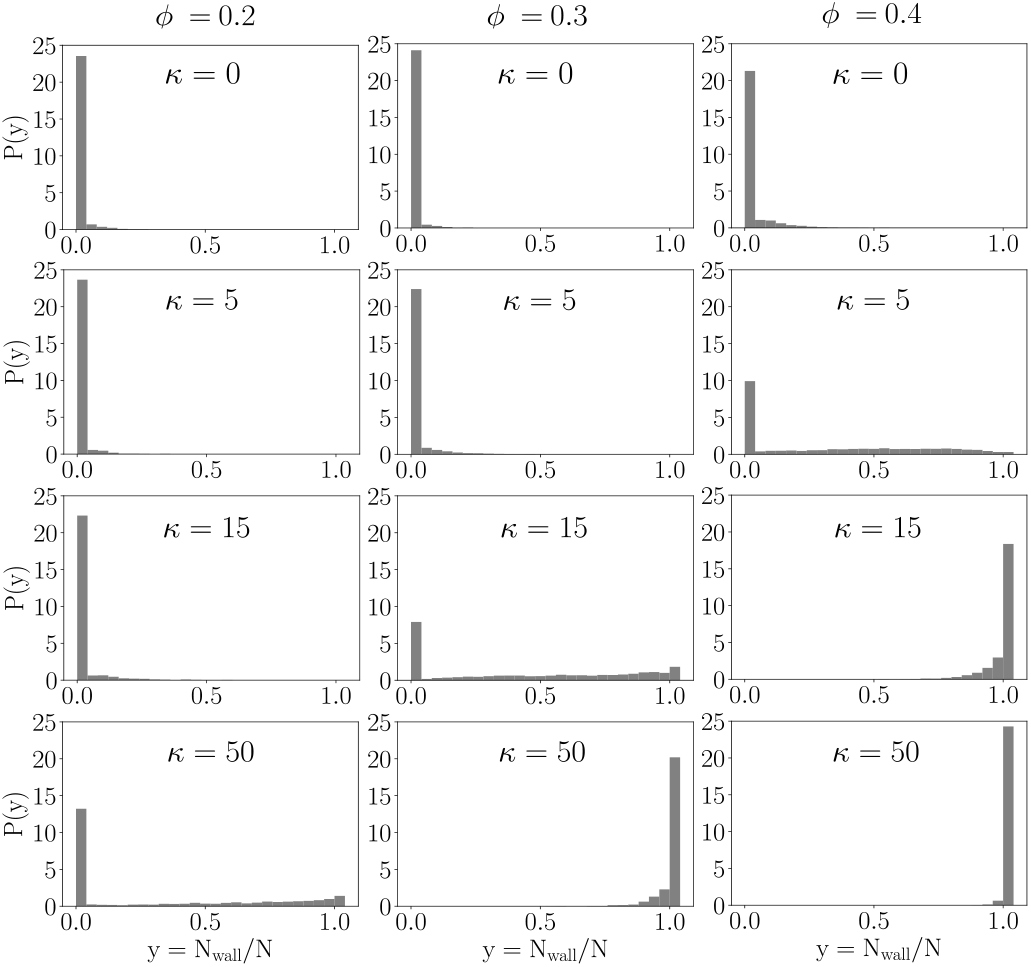
Probability density of the fraction of monomers near a confining wall. The volume fraction of crowding particles (*ϕ* = 0.2, 0.3, 0.4) increases from left to right. The bending stiffness of the polymer (*κ* = 0, 5, 15, 50) increases from top to bottom. Simulations were carried out with σ_*c*_ = 0.8σ_*m*_.

Similar behavior was observed for *ϕ* = 0.3 and *κ* = 15, in which the adsorbed polymer adopted conformations with trains and tails but no loops. However, a greater degree of flexibility is evident in the tails compared with the previous case. For *ϕ* = 0.4 and *κ* = 5, the partially adsorbed polymer formed trains, tails and loops. In this regime, the polymer is sufficiently flexible so that the energetic penalty associated with forming loops is offset by entropic gains.

The effects of crowding on the polymer are further illuminated by the radius of gyration of the polymer. Figure 4a shows the radius of gyration as a function of bending stiffness in uncrowded (*ϕ* = 0) conditions. Figures 4b and 4c show, for different *κ*, how the radius of gyration varies as a function of the crowding fraction (*R_g_*(*ϕ*)) in relation to the corresponding value in uncrowded conditions (*R_g_*(0)). For the purely flexible polymer chain (*κ* = 0), the radius of gyration decreases with an increase in crowding, which is consistent with previous work on flexible polymers.^15^ The decrease is less pronounced for the semiflexible polymer with *κ* = 5 in the regime without adsorption (*ϕ* ≤ 0.3), and no decrease is evident for stiffer polymers. A notable feature is that at sufficiently large values of *ϕ*, there is an increase in *R_g_*. The largest relative increase in size is observed for intermediate values of the stiffness (*κ* = 10 and 15), with a nearly 20% increase in *R_g_* at *ϕ* = 0.4. These increases occur in regimes in which the polymer adsorbs to the wall and flattens against it. The increase is considerably smaller for *κ* = 50 and especially *κ* = 100 because these polymers are relatively extended even in the bulk.

**FIG. 3.**
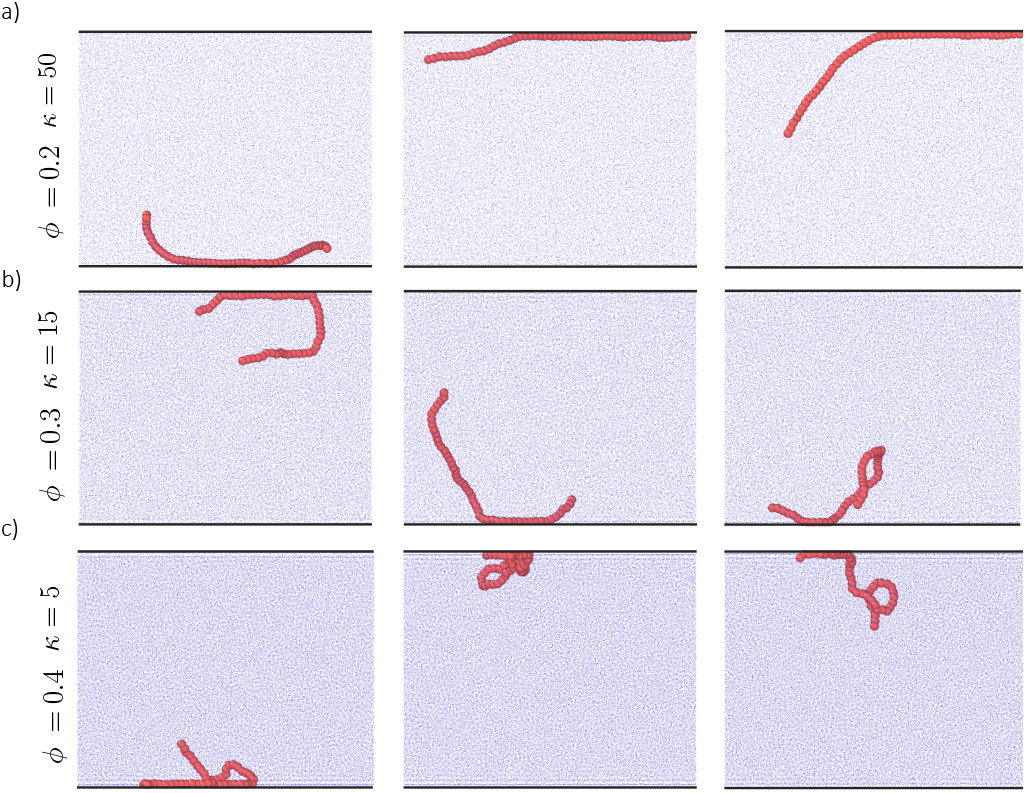
Snapshots of partially adsorbed semiflexible polymers. Boundary conditions are periodic in the *x*- and *y*-directions; confining walls in the *z*-direction are shown by black lines. The size of the crowders is σ_*c*_ = 0.8σ_*m*_. (a) - (c) represent three cases (different values of *κ* and *ϕ*) that exhibit partial adsorption of the polymer. Three snapshots are shown for each case. The system is viewed along the *y*-axis, and the crowding particles are minimized in the visualization to more clearly reveal the polymer.

To further characterize the shape of the polymer, we calculated the radius of gyration parallel to the wall (*R*_*g*,||_) and perpendicular to the wall (*R*_*g*,⊥_), using

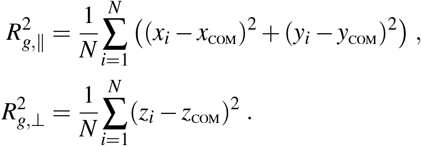

Figure 5 shows the average values of *R*_*g*,||_ and *R*_*g*,⊥_ as a function of the crowding fraction. For the flexible polymer (*κ* = 0), the behavior of each reflects the decrease seen in *R_g_*, indicating that crowding leads to a compaction in directions both parallel and perpendicular to the wall.

For *κ* ≥ 5, *R*_*g*,||_ increases when the polymer is adsorbed to the wall. Small increases are seen for partially adsorbed polymers (e.g., *κ* = 5 and *ϕ* = 0.4). Larger increases are seen for fully adsorbed polymers, although the relative increase in *R*_*g*,||_ is less for large values of *κ*. When *R*_*g*,||_ increases, there is a concomitant decrease in *R*_*g*⊥_, with the value approaching zero for strongly adsorbed polymers. These results are consistent with the polymer flattening against the wall (decreasing *R*_*g*⊥_) and increasing its size in the directions parallel to the wall (increasing *R*_*g*,||_). The relative increase in the parallel direction is largest for smaller values of *κ* because the polymer can adopt more compact conformations when not adsorbed. In contrast, in the limit of a rigid-rod polymer (*κ* → ∞), *R_g_* would remain constant upon adsorption and changes in *R*_*g*⊥_ and *R*_*g*,||_ would reflect the change in the orientational degrees of freedom only.

**FIG. 4.**
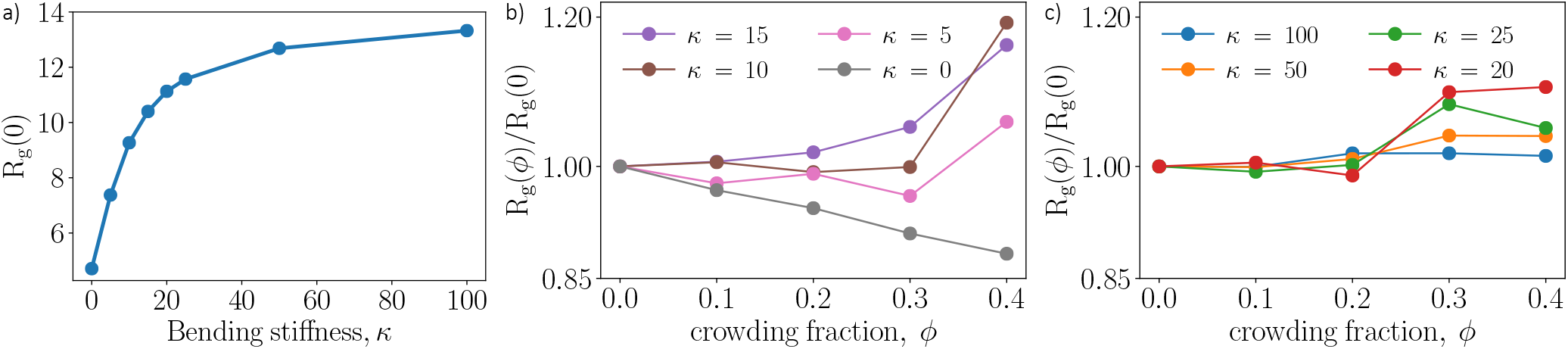
(a) Mean radius of gyration of the polymer in uncrowded conditions (*R_g_*(0)) as a function of bending stiffness (*κ*). (b),(c) Mean radius of gyration at crowding fraction ϕ scaled by the corresponding value in uncrowded conditions (*R_g_* (*ϕ*)/*R_g_*(0)). For clarity, lower and higher values of the bending stiffness are shown separately. The size of the crowders is σ_*c*_ = 0.8σ_*m*_.

**FIG. 5.**
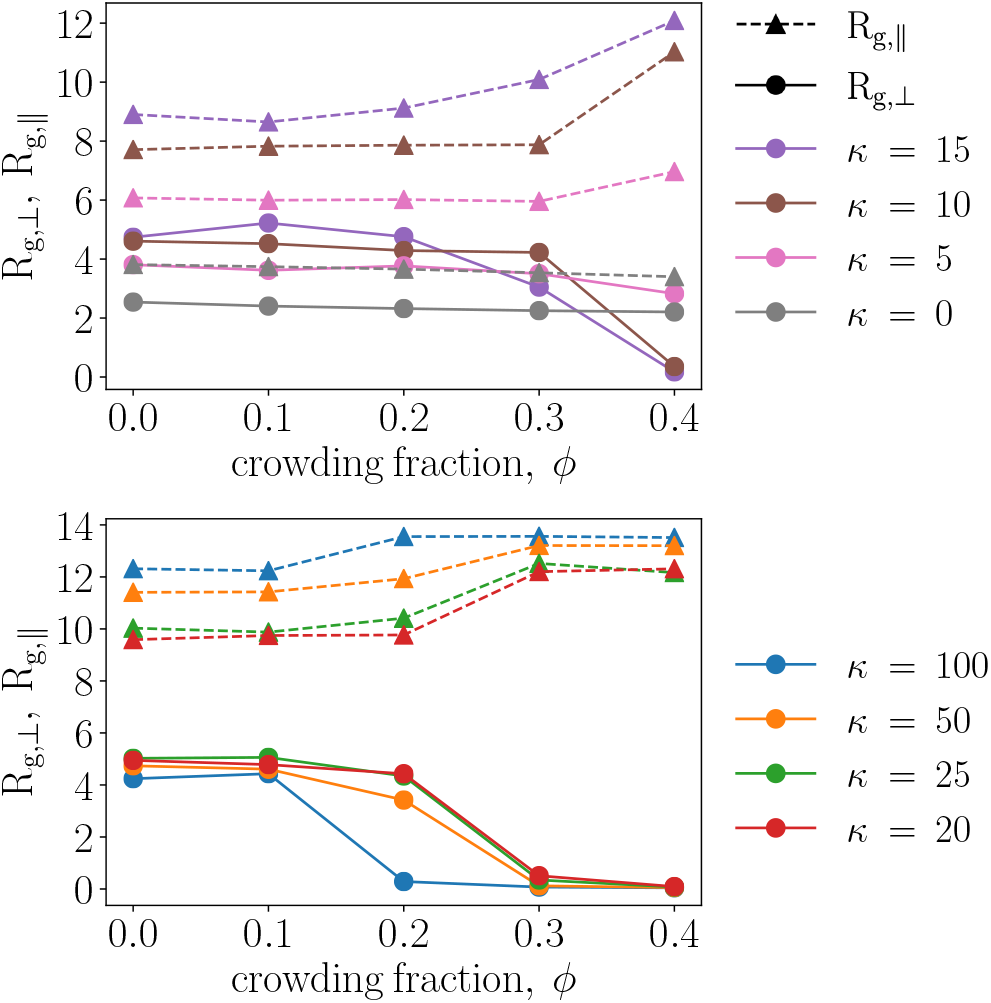
Mean radius of gyration parallel to the wall (*R*_*g*,||_, dashed) and perpendicular to the wall (*R*_*g*⊥_, solid), for various values of the bending stiffness (*κ*).

### C. Adsorption in spherical confinement

Confining semiflexible polymers in small regions can significantly impact their conformations,^46^ but even modestly confined systems can have an impact because the bending energy of the adsorbed polymer, imposed by curvature of the surface, can compete with depletion interactions. We carried out simulations of a semiflexible polymer with crowding in a spherical domain with a diameter of 60σ. For comparison, the volume is about 13% larger than that of the cuboid in the previous sections, and the ratio of the surface area to volume is twice as large.

Figure 6 shows the average fraction of monomers near the spherically confining wall as a function of *κ* for various crowding fractions. Like the previous case with flat walls, contact of the polymer with the wall is promoted by larger values of both the crowding fraction and bending stiffness. In contrast with the previous case, the transition to strong adsorption occurs at smaller values of the bending stiffness. This is evident in the regions in which the polymer is partially adsorbed, where the fraction of monomers near the wall is larger for the spherically confined case than for case with flat walls. These differences are likely primarily because of the increased surface area to volume ratio, and hence the larger effective volume near the wall in the spherically confined case.

**FIG. 6.**
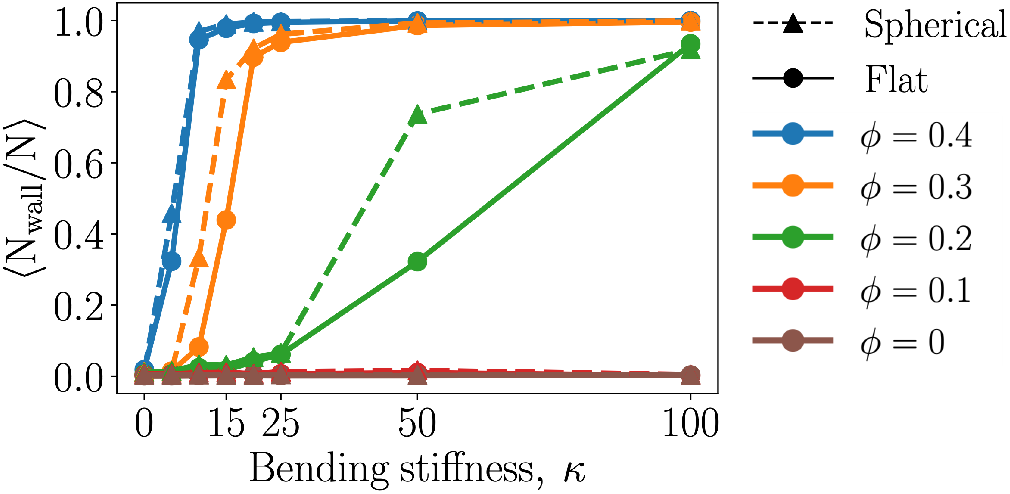
Average fraction of monomers near a wall as a function of the bending stiffness. Solid lines correspond to a flat wall and dashed lines correspond to spherical confinement with a radius of 30σ. The size of the crowders is σ_*c*_ = 0.8σ_*m*_.

**FIG. 7.**
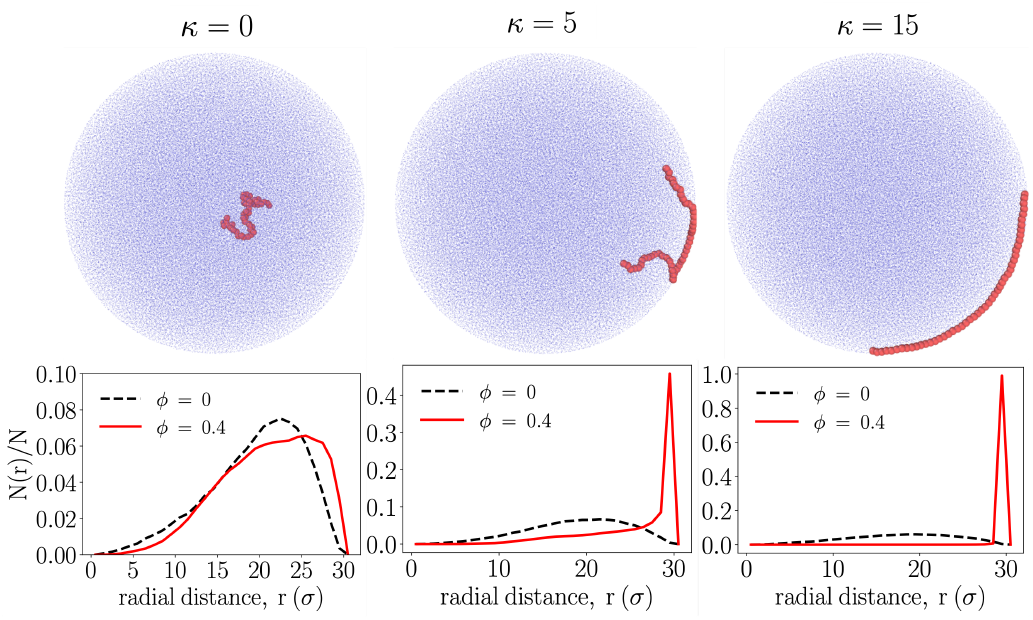
Snapshots of the polymer in spherical confinement with *ϕ* = 0.4. Three values of the bending stiffness are shown. The plots below correspond to the same values of *κ* and show the average fraction of monomers, *N*(*r*)/*N*, as a function of the radial distance, r. For each, results are shown for *ϕ* = 0 and 0.4.

Figure 7 shows representative snapshots of the polymer in spherical confinement for different values of *κ* at *ϕ* = 0.4. There is no adsorption at *κ* = 0, partial adsorption at *κ* = 5, and strong adsorption at *κ* = 15. The partially adsorbed polymer has a train with two tails that extend into the interior of the sphere. The strongly adsorbed polymer follows the contour of the sphere. The plots in Fig. 7 show the corresponding time-averaged distribution of the fraction of polymer beads as a function of the radial distance, r. For comparison, the results for the uncrowded system are also shown. For the uncrowded system, the fraction of monomers first increases with increasing radial distance because the volume grows with increasing *r* (∝ *r*^2^*dr*). The fraction of monomers decreases close to the wall because there is an effective repulsion between the wall and the center of mass of the polymer that arises due to the wall restricting conformations of the polymer. For *ϕ* = 0.4, the polymer does not adsorb to the wall at *κ* = 0, but the distribution is shifted slightly toward larger values of *r*. This indicates that the polymer resides slightly closer to the wall on average. The effect of crowding is likely to make the effective interaction between the polymer and wall less repulsive.^17^ A strong enhancement of the bead density is observed near the wall for *κ* = 5 (partial adsorption) and *κ* = 15 (strong adsorption). Like before, the adsorption of stiffer polymers is promoted because they lose less conformational entropy than their flexible counterpart upon adsorption.

The adsorption of a semiflexible polymer to a flat surface is promoted by increasing *κ*. Adsorption to a curved surface, however, introduces an additional competing energy contribution: the energetic penalty associated with bending an adsorbed semiflexible polymer along the contour of the surface. This suggests that adsorption should be suppressed at sufficiently large values of the bending stiffness.

To explore this, we carried out simulations at larger values of *κ* for the polymer in spherical confinement. Figure 8 shows the fraction of polymer beads near the wall for bending stiffness up to *κ* = 2000. There was no adsorption for *ϕ* ≤ 0.1 over the extended range of *κ*, but there were regimes of strong adsorption for *ϕ* = 0.2, 0.3, and 0.4. For *ϕ* = 0.2, this regime emerged for *κ* > 100. However, when *κ* was sufficiently large (*κ* = 2000), we observed another regime in which the polymer was not adsorbed. In contrast, the cases with *ϕ* = 0.3 and 0.4 remained strongly adsorbed at *κ* = 2000. The case with *ϕ* = 0.2 generates the weakest depletion interactions of the three cases. The non-adsorbed regime at high *κ* emerged due to the bending energy penalty imposed by being in close proximity to the curved wall. At *ϕ* = 0. 3 and 0. 4, the depletion interactions were still sufficiently strong to promote adsorption, highlighting the interplay between crowding-induced depletion interactions and curvature of the surface.

**FIG. 8.**
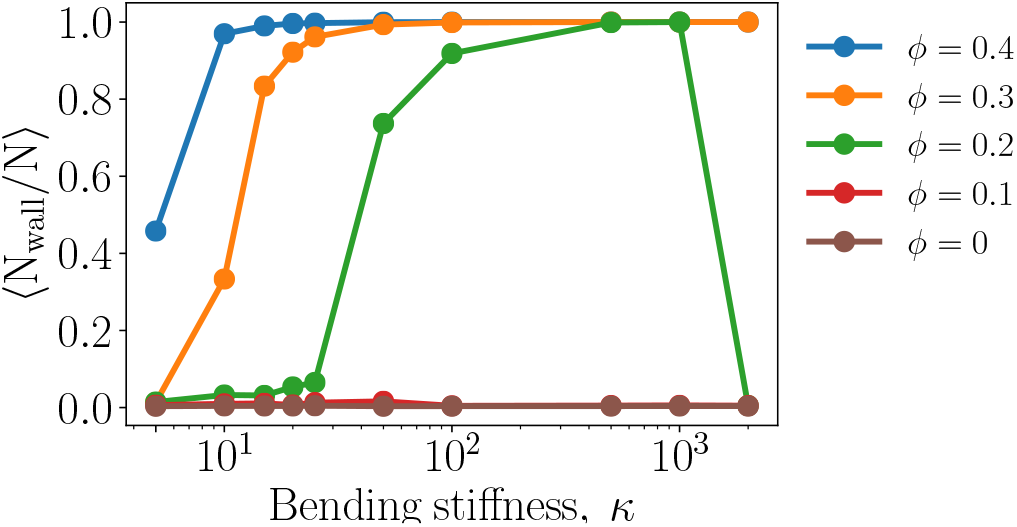
Average fraction of monomers near the wall in spherical confinement. The range of *κ* is larger than in previous figures.

## IV. DISCUSSION

The adsorption of semiflexible polymers is a problem of fundamental interest that is relevant to cellular and cell-free biological systems, where macromolecular crowding can lead to depletion interactions that affect biopolymers such as DNA and actin. In this work, we used computer simulations to study the crowding-induced adsorption of a bead-spring model of a semiflexible polymer. Unlike most previous works on the adsorption of semiflexible polymers, we explicitly accounted for crowding particles, which were modeled as purely repulsive particles.

Figure 1 demonstrates that polymer adsorption is promoted by stiffer polymers, smaller crowding particles, and larger volume fractions of crowders. With increasing bending stiffness (*κ*), we observed transitions from non-adsorbed states to partially and then fully adsorbed states. The transitions occurred at smaller values of *κ* as the volume fraction of crowders (*ϕ*) increased. Smaller crowding particles enhanced the effective polymer-wall attraction at the same volume fraction, shifting the adsorption transition to smaller values of the polymer stiffness. The transitions were accompanied by changes in the radius of gyration (Figs. 4 and 5) demonstrating that the polymer flattened against the wall while increasing in size in directions parallel to the wall.

Physically, adsorption of a polymer due to crowding results from an increase in the entropy of the crowders (due to an increase in the accessible volume) that exceeds the loss of entropy associated with the polymer. Semiflexible polymers are more likely than flexible polymers to adsorb to a wall because the conformational entropy of a polymer depends on its persistence length and stiffer polymers experience a smaller loss of conformational entropy upon adsorption.^22,31–33^ Additionally, the strength of depletion interactions increases with increasing concentration of depletants.^1^ Our results showed that larger values of *κ* and *ϕ* promoted polymer adsorption, which is consistent with these physical underpinnings of depletion interactions and polymer adsorption.

For partially adsorbed polymers, the bending rigidity of the polymer impacted the conformations of the polymer. Loops were observed when partial adsorption occurred at smaller values of *κ*, but not when the polymer was partially adsorbed at larger values. In this regime, only one or both ends of the polymer were not in contact with the wall. Finally, we showed that curved surfaces, such as those observed in cells and vesicles, can give rise to behavior in which the number of monomers in contact changes nonmonotonically with *κ*, indicating a desorption transition at large *κ*. In this regime, the energetic cost of bending the polymer competes with the crowding- and stiffness-dependent depletion interactions.

In this work, we considered monodisperse crowding particles and static walls. In reality, cellular environments are crowded with macromolecules of different shapes and sizes, which can can give rise to unexpected effects on the shapes of semiflexible polymers^19^ and can affect the depletion interaction experienced by polymers.^47,48^ Additionally, cell membranes are deformable, and adsorption of a semiflexible polymer can lead to an interplay between the shapes of the membrane and polymer due to competing bending energies.^36^ In the future, it would be interesting to study the effect of crowder polydispersity and membrane flexibility on the adsorption of semiflexible polymers in crowded environments.

The cellular environment is replete with biopolymers that have a variety of persistence lengths. Our work sheds light on how crowding and the presence of surfaces – both features of cells – affect the spatial organization of biopolymers through their interactions with surfaces. In the context of cell-free systems, our work provides guidance on how crowding can be used to rationally organize components with different persistence lengths.

## V. DATA AVAILABILITY

The data that support the findings of this study are available from the corresponding author upon request.

## ACKNOWLEDGMENTS

This research was conducted as part of the Interface Directed Assembly Theme at the Center for Nanophase Materials Sciences, which is a DOE Office of Science User Facility.

## Notes

### Competing Interest Statement

The authors have declared no competing interest.

## References

1 S. Asakura and F. Oosawa, “On interaction between two bodies immersed in a solution of macromolecules,” J. Chem. Phys. 22, 1255–1256 (1954).

2 K. Binder, P. Virnau, and A. Statt, “Perspective: The Asakura Oosawa model: A colloid prototype for bulk and interfacial phase behavior,” J. Chem. Phys. 141, 559 (2014).

3 R. Tuinier and H. N. Lekkerkerker, Colloids and the Depletion Interaction(Springer Netherlands, 2011).

4 D. Marenduzzo, K. Finan, and P. R. Cook, “The depletion attraction: an underappreciated force driving cellular organization,” J. Cell. Biol. 175, 681–686 (2006).

5 D. Marenduzzo, C. Micheletti, and P. R. Cook, “Entropy-driven genome organization,” Biophys. J. 90, 3712–3721 (2006).

6 J. Pelletier, K. Halvorsen, B.-Y. Ha, R. Paparcone, S. J. Sandler, C. L. Woldringh, W. P. Wong, and S. Jun, “Physical manipulation of the Escherichia coli chromosome reveals its soft nature,” Proc. Natl. Acad. Sci. U.S.A 109, E2649–E2656 (2012).

7 O. L. Kantidze and S. V. Razin, “Weak interactions in higher-order chromatin organization,” Nucleic Acids Res. 48, 4614–4626 (2020).

8 C. Jeon, Y. Jung, and B.-Y. Ha, “A ring-polymer model shows how macromolecular crowding controls chromosome-arm organization in Escherichia coli,” Sci. Rep. 7, 1–10 (2017).

9 A. Papantonis and P. R. Cook, “Transcription factories: genome organization and gene regulation,” Chem. Rev. 113, 8683–8705 (2013).

10 H.-X. Zhou, G. Rivas, and A. P. Minton, “Macromolecular crowding and confinement: biochemical, biophysical, and potential physiological consequences,” Annu. Rev. Biophys. 37, 375–397 (2008).

11 S. E. Norred, P. M. Caveney, G. Chauhan, L. K. Collier, C. P. Collier, S. M. Abel, and M. L. Simpson, “Macromolecular crowding induces spatial correlations that control gene expression bursting patterns,” ACS Synth. Biol. 7, 1251–1258 (2018).

12 M. Delarue, G. P. Brittingham, S. Pfeffer, I. Surovtsev, S. Pinglay, K. Kennedy, M. Schaffer, J. Gutierrez, D. Sang, G. Poterewicz, et al., “mTORC1 controls phase separation and the biophysical properties of the cytoplasm by tuning crowding,” Cell 174, 338–349 (2018).

13 T. Kaur, I. Alshareedah, W. Wang, J. Ngo, M. M. Moosa, and P. R. Banerjee, “Molecular crowding tunes material states of ribonucleoprotein condensates,” Biomolecules 9, 71 (2019).

14 R. J. Ellis, “Macromolecular crowding: an important but neglected aspect of the intracellular environment,” Curr. Opin. Struct. Biol. 11, 114–119 (2001).

15 H. Kang, P. A. Pincus, C. Hyeon, and D. Thirumalai, “Effects of macromolecular crowding on the collapse of biopolymers,” Phys. Rev. Lett. 114, 068303 (2015).

16 T. N. Shendruk, M. Bertrand, H. W. de Haan, J. L. Harden, and G. W. Slater, “Simulating the entropic collapse of coarse-grained chromosomes,” Biophys. J. 108, 810–820 (2015).

17 G. Chauhan, M. L. Simpson, and S. M. Abel, “Crowding-induced interactions of ring polymers,” Soft Matter 17, 16–23 (2021).

18 A. Lau, A. Prasad, and Z. Dogic, “Condensation of isolated semi-flexible filaments driven by depletion interactions,”Europhys. Lett. 87, 48006 (2009).

19 H. Kang, N. M. Toan, C. Hyeon, and D. Thirumalai, “Unexpected swelling of stiff DNA in a polydisperse crowded environment,” J. Am. Chem. Soc. 137, 10970–10978 (2015).

20 S. E. Norred, R. M. Dabbs, G. Chauhan, P. M. Caveney, C. P. Collier, S. M. Abel, and M. L. Simpson, “Synergistic interactions between confinement and macromolecular crowding spatially order transcription and translation in cell-free expression,” bioRxiv, 445544 (2018).

21 N. Biswas, M. Ichikawa, A. Datta, Y. T. Sato, M. Yanagisawa, and K. Yoshikawa, “Phase separation in crowded micro-spheroids: DNA-PEG system,” Chem. Phys. Lett. 539, 157–162 (2012).

22 D. Welch, M. Lettinga, M. Ripoll, Z. Dogic, and G. A. Vliegenthart, “Trains, tails and loops of partially adsorbed semi-flexible filaments,” Soft Matter 11, 7507–7514 (2015).

23 T. Hoppe and A. P. Minton, “An equilibrium model for the combined effect of macromolecular crowding and surface adsorption on the formation of linear protein fibrils,” Biophys. J. 108, 957–966 (2015).

24 A. P. Minton, “The cumulative effect of surface adsorption and excluded volume in 2D and 3D on protein fibrillation,” Biophys. J. 117, 1666–1673 (2019).

25 P. De Gennes, “Polymers at an interface; a simplified view,” Adv. Colloid Interface Sci. 27, 189–209 (1987).

26 R. R. Netz and D. Andelman, “Neutral and charged polymers at interfaces,” Phys. Rep. 380, 1–95 (2003).

27 J. Baschnagel, H. Meyer, J. Wittmer, I. Kulić, H. Mohrbach, F. Ziebert, G.-M. Nam, N.-K. Lee, and A. Johner, “Semiflexible chains at surfaces: Worm-like chains and beyond,” Polymers 8, 286 (2016).

28 R. Simha, H. Frisch, and F. Eirich, “The adsorption of flexible macromolecules,” J. Phys. Chem. 57, 584–589 (1953).

29 A. Milchev and K. Binder, “Static and dynamic properties of adsorbed chains at surfaces: Monte carlo simulation of a bead-spring model,” Macromolecules 29, 343–354 (1996).

30 E. Eisenriegler, K. Kremer, and K. Binder, “Adsorption of polymer chains at surfaces: Scaling and monte carlo analyses,” J. Chem. Phys. 77, 6296–6320 (1982).

31 A. Milchev and K. Binder, “Linear dimensions of adsorbed semiflexible polymers: What can be learned about their persistence length?” Phys. Rev. Lett. 123, 128003 (2019).

32 T. Sintes, K. Sumithra, and E. Straube, “Adsorption of semiflexible polymers on flat, homogeneous surfaces,” Macromolecules 34, 1352–1357 (2001).

33 J. Jiang, “Nonmonotonic adsorption behavior of semiflexible polymers,” J. Chem. Phys. 153, 034902 (2020).

34 T. A. Kampmann, H.-H. Boltz, and J. Kierfeld, “Controlling adsorption of semiflexible polymers on planar and curved substrates,” J. Chem. Phys. 139, 034903 (2013).

35 W. Chien and Y.-L. Chen, “Confinement, curvature, and attractive interaction effects on polymer surface adsorption,” J. Chem. Phys. 147, 064901 (2017).

36 B. Li and S. M. Abel, “Shaping membrane vesicles by adsorption of a semiflexible polymer,” Soft Matter 14, 185–193 (2018).

37 X. Zhou, F. Guo, K. Li, L. He, and L. Zhang, “Entropy-induced separation of binary semiflexible ring polymer mixtures in spherical confinement,” Polymers 11, 1992 (2019).

38 J.-Z. Zhang, X.-Y. Peng, S. Liu, B.-P. Jiang, S.-C. Ji, and X.-C. Shen, “The persistence length of semiflexible polymers in lattice Monte Carlo simulations,” Polymers 11, 295 (2019).

39 P. Gutjahr, R. Lipowsky, and J. Kierfeld, “Persistence length of semiflexible polymers and bending rigidity renormalization,” Europhys. Lett. 76, 994 (2006).

40 J. D. Weeks, D. Chandler, and H. C. Andersen, “Role of repulsive forces in determining the equilibrium structure of simple liquids,” J. Chem. Phys. 54, 5237–5247 (1971).

41 S. Plimpton, “Fast parallel algorithms for short-range molecular dynamics,” Tech. Rep. (Sandia National Labs., Albuquerque, NM (United States), 1993).

42 S. Plimpton, A. Thompson, P. Crozier, and A. Kohlmeyer, “LAMMPS molecular dynamics simulator,” URL http://lammps.sandia.gov (2011).

43 A. Stukowski, “Visualization and analysis of atomistic simulation data with OVITO–the open visualization tool,” Model. Simul. Mater. Sci. Eng. 18, 015012 (2009).

44 J. K. Armstrong, R. B. Wenby, H. J. Meiselman, and T. C. Fisher, “The hydrodynamic radii of macromolecules and their effect on red blood cell aggregation,” Biophys. J. 87, 4259–4270 (2004).

45 L. Dai, J. J. Jones, J. R. van der Maarel, and P. S. Doyle, “A systematic study of DNA conformation in slitlike confinement,” Soft Matter 8, 2972–2982 (2012).

46 S. Mirzaeifard and S. M. Abel, “Confined semiflexible polymers suppress fluctuations of soft membrane tubes,” Soft Matter 12, 1783–1790 (2016).

47 A. Chen and N. Zhao, “Comparative study of the crowding-induced collapse effect in hard-sphere, flexible polymer and rod-like polymer systems,” Phys. Chem. Chem. Phys. 21, 12335–12345 (2019).

48 W. M. Mardoum, S. M. Gorczyca, K. E. Regan, T.-C. Wu, and R. M. Robertson-Anderson, “Crowding induces entropically-driven changes to DNA dynamics that depend on crowder structure and ionic conditions,” Front. Phys. 6, 53 (2018).

